# How lizards fly: A novel type of wing in animals

**DOI:** 10.1101/086496

**Authors:** J. Maximilian Dehling

## Abstract

Flying lizards of the genus *Draco* are famous for their gliding ability, using an aerofoil formed by winglike patagial membranes and supported by elongated thoracic ribs. It has remained unknown, however, how the lizards manoeuvre during flight. Here, I show that the patagium is deliberately grasped and controlled by the forelimbs while airborne. This type of composite wing is unique inasmuch as the lift-generating and the controlling units are formed independently by different parts of the body and are connected to each other only for the duration of the flight. The major advantage for the lizards is that the forelimbs keep their entire movement range and functionality for climbing and running when they are not used as the controlling unit of the wing. These findings not only shed a new light on the flight of *Draco* lizards but also have implications for the interpretation of gliding performance in fossil species.

## INTRODUCTION

A number of vertebrates as well as invertebrates are known to perform gliding flights (Dudley *et al.*, 2007; Socha *et al.*, 2015; Lingham-Soliar, 2015). Flying Lizards of the agamid genus *Draco* are the most specialized and best-studied gliding reptiles (McGuire, 2003; McGuire & Dudley, 2005, 2011; Socha *et al.*, 2015). Their patagium is supported by five to seven greatly elongated thoracic ribs and spread by specialized iliocostalis and intercostal muscles (Colbert, 1967; Russell & Dijkstra, 2001; McGuire, 2003; Dudley *et al.*, 2007; McGuire & Dudley, 2011; Lingham-Soliar, 2015). It is commonly assumed that flying lizards use the unfurled patagium to glide but hold the forelimbs free in front of the body while airborne. This assumption was manifested about 300 years ago, when the first preserved specimens were brought to Europe and reports on Flying Lizards were accompanied with drawings showing artistic interpretations of “gliding” lizards holding their forelimbs in front of the body (e.g. Seba, 1734; Marsden, 1811; Günther, 1872; Maindron, 1890). The patagium-associated musculature has been suspected to control the direction of the glide path (Colbert, 1967; Russell & Dijkstra, 2001; Dudley *et al.*, 2007; McGuire & Dudley, 2011; Lingham-Soliar, 2015), but it has remained unclear how the lizards are able to manoeuvre in the air (McGuire & Dudley 2011).

Anatomical properties of the patagium as well as behavioural observations challenge the assumption that the associated muscles alone can perform the sophisticated movements required for manoeuvring. (1) Only two muscles insert the first elongated ribs (Colbert, 1967) and therefore allow movements in only a limited number of different directions, mainly spreading (forward) and furling (backward) the patagium. (2) The patagium-spreading muscles stem from musculature originally used for breathing (Colbert, 1967; John, 1970). In the original state, the intercostal muscles of both sides contract simultaneously in order to expand and contract the thorax (e.g. Ratnovsky *et al.*, 2008). If the muscles of only one side contracted, the thorax would be rotated. One-sided contractions of intercostal muscles could so far be demonstrated only in anaesthesized dogs and in humans in response to passive rotations of the thorax (Decramer *et al.*, 1986; Whitelaw *et al.*, 1992), and therefore it seems unlikely that *Draco* lizards are able to deliberately execute one-sided contractions of these muscles. (3) The patagium is spread not only to form an aerofoil but also for display in intraspecific communication, and photographs and observations of display in different species of *Draco* indicate that both sides of the patagium are always moved simultaneously (Hairstone, 1957; John, 1967; Mori & Hikida, 1993; McGuire & Dudley, 2011; J. M. Dehling, unpubl. data). If *Draco* lizards were able to perform sophisticated one-sided movements required for glide-path control with their patagium-associated muscles alone, they would probably show them during display as well, but all they display are simultaneous spreading and furling of the patagium on both sides. Therefore, it appears unlikely that *Draco* lizards are able to manoeuvre in the air using the specialized muscles of the trunk alone.

Here, I report on the results of a study I carried out on the aerial behaviour of Dussumier's Flying Lizard (*Draco dussumieri*) in order to investigate if the patagium is controlled in a different way. Observations and documentations of the gliding flight in the habitat are supplemented with examinations of morphological characteristics in preserved specimens of *D. dussumieri* and 17 other species of the genus *Draco*. My findings demonstrate that the patagium is actually controlled by the forelimbs and thus reveal a hitherto unknown type of wing in animals.

## MATERIAL AND METHODS

### Behavioural observations

I observed gliding flights of *Draco dussumieri* in an abandoned areca nut (*Areca catechu*) plantation near the town of Agumbe, southwestern India (13.517628°N, 75.088542°E, WGS 84), during the late morning and early afternoon (10–12 h, 13–14.30 h) on seven non-consecutive days in March 2015. The observations were made in a non-experimental approach on the natural behaviour in the habitat, where the lizards performed gliding flights from one tree to another. No animal was captured, handled, or manipulated in any other way during the study. A total of approximately 200 gliding flights performed by at least seven different individuals were observed, partly using Minox 10x50 binoculars. I documented about 50 glides photographically, focusing mainly on the initial phases of the gliding flight. Sequential short-exposure photographs were taken at a rate of 5.5 frames per second with a Nikon D600 full-frame digital single-lens reflex camera equipped with a Nikon AF-S 200–400 mm telephoto zoom lens (manually focused).

### Morphological examination

In order to corroborate observations of morphological adaptations to the gliding flight in *Draco* lizards, I examined voucher specimens of 18 species of *Draco* and 21 species of 12 representative genera of other arboreal Asian agamid lizards deposited in the herpetological collection of the Zoologisches Forschungsmuseum Alexander Koenig (ZFMK), Bonn, Germany (online supporting information, Table S1). I took measurements (to the nearest 0.1 mm) with a digital calliper of snout-vent length (SVL, from tip of snout to vent), arm length (AL, from the forelimb insertion to the distal end of the antebrachium, measured with the arm extended perpendicularly to the median body plane), and length of the leading edge of the patagium (LL, from the insertion of the first elongated rib to the point where the leading edge starts to bend posteriorly; given as a percentage of the corresponding arm length, rounded to the nearest 1 %). I checked the ability to deviate the wrist ulnarly and radially in all specimens. The results of the examination are given in the online supporting information, Table S1.

## RESULTS

Sequential photographs of gliding *Draco* lizards revealed that the patagial musculature is not the major element controlling the glide path. The explanation of how *Draco* lizards achieve manoeuvrability while airborne is surprising: Instead of being held free in front of the body, as previously assumed, the powerful and highly movable forelimbs are attached to the leading edge of the patagium for the duration of the flight and control the aerofoil (Fig. 1).

**Figure 1.**
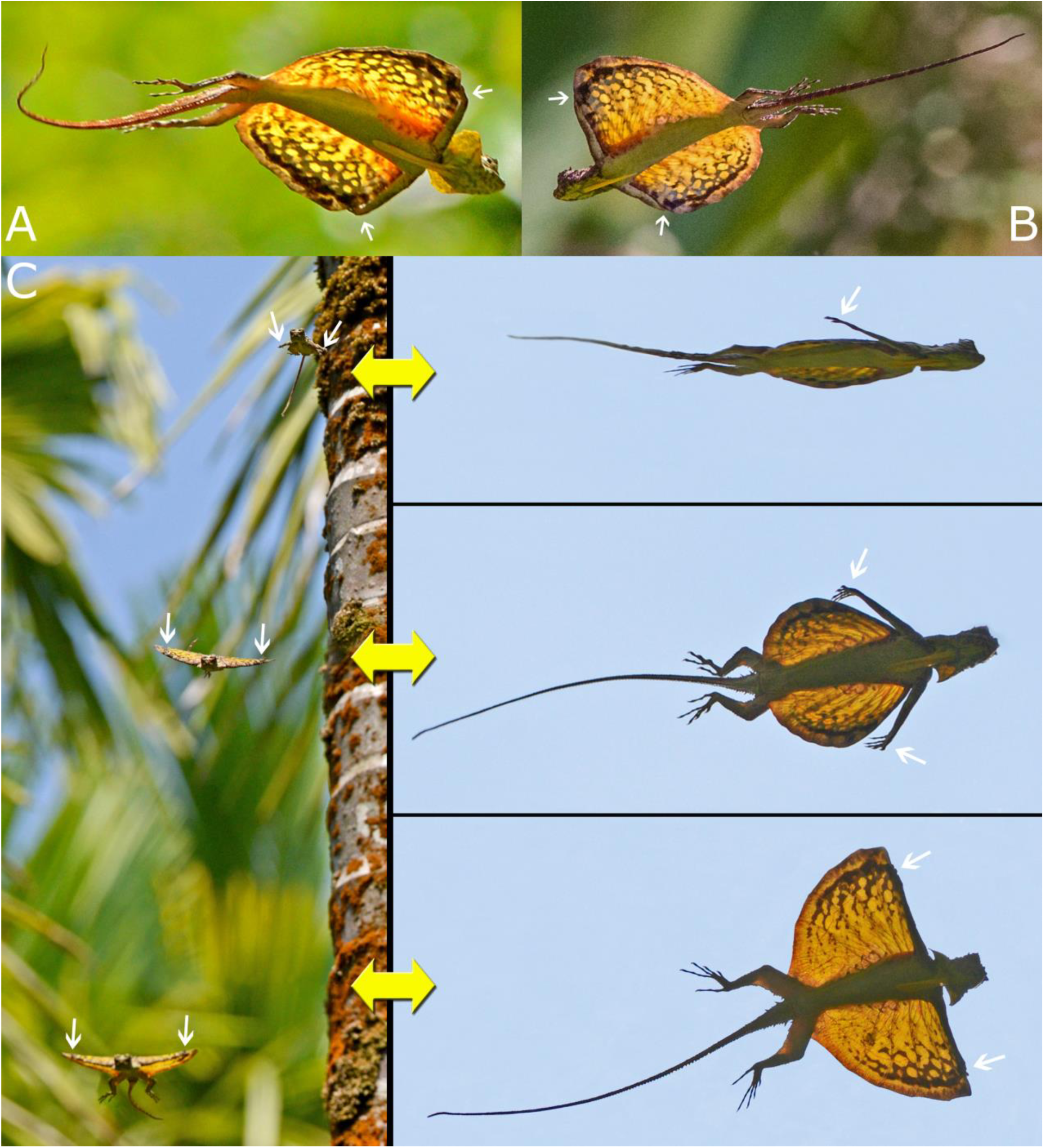
Gliding specimens of *Draco dussumieri*. A, B, in mid-flight; note the attachment of the arms to the leading edge of the patagium and the marked adduction of the wrists. C, formation of the composite wing during the initial phases of the gliding flight of *Draco dussumieri* seen from the front (left) and from below (right; corresponding photos of the same phases). The lizard jumps from the tree, reorients the body dorsoventrally and starts to spread the anterior ribs; the extended arms reach behind the back (top). The anterior ribs are further spread by the trunk musculature; the hands grasp the leading edge of the patagium and pull it forward (middle). The patagium is fully extended and controlled by the forelimbs; the glide path becomes more horizontal (bottom). White arrows indicate the positions of the hands.

The movements and actions of the forelimbs and the patagium followed a certain pattern in all observed and documented gliding flights. Initially, a lizard launched itself from the tree with a jump and descended head first. After takeoff, it reoriented its body dorsoventrally, the extended forelimbs reached behind the back, and the trunk muscles started to spread the anterior patagium-supporting ribs (Fig. 1C). When the hands got hold of the outer margins of the leading edge of the patagium, they pulled it forward, and the patagium was unfurled to its full extent (Fig. 1C). The flight path then soon became more horizontal. The forelimbs remained attached to the patagium for the duration of the gliding flight (Fig. 1A, B), enabling changes of direction through unilateral movements of the aerofoil. Shortly before reaching the landing point, the leading edge of the patagium was raised above the body plane and the hind limbs were lowered, causing a change of the angle of attack and an upturn of the glide path. Immediately before landing, the forelimbs released the patagium and were flexed forward to diminish the impact and take hold of the surface. During the landing process, the patagium was furled against the sides of the body.

During the gliding flight, the fully extended patagium was strongly cambered, and the lizards actively arched their backs when airborne and thereby increased the camber of the aerofoil (Fig. 2). The attached forelimbs formed a straight, thick leading edge of the aerofoil compared to the thin trailing edge (Figs. 1, 2). When pulling the patagium forward and holding on to it, the wrist was deviated ulnarly about 90° to the extended arm (Figs. 1, 3).

**Figure 2.**
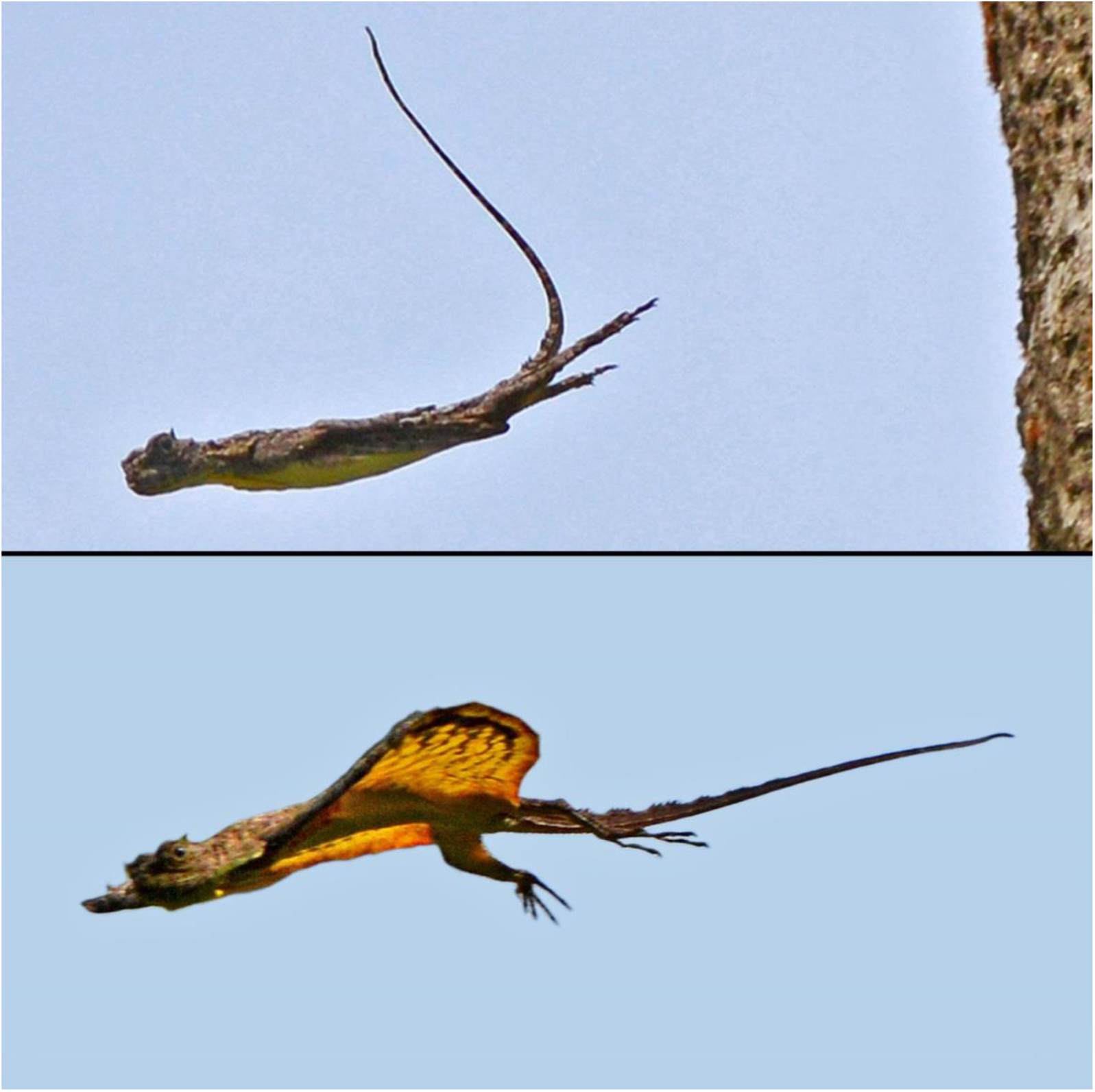
*Draco dussumieri* during takeoff jump (top) and during gliding flight (bottom). Note the cambered shape of the patagium and the arching of the back when the patagium is extended (bottom), in comparison to the straight back during takeoff with furled patagium (top).

**Figure 3.**
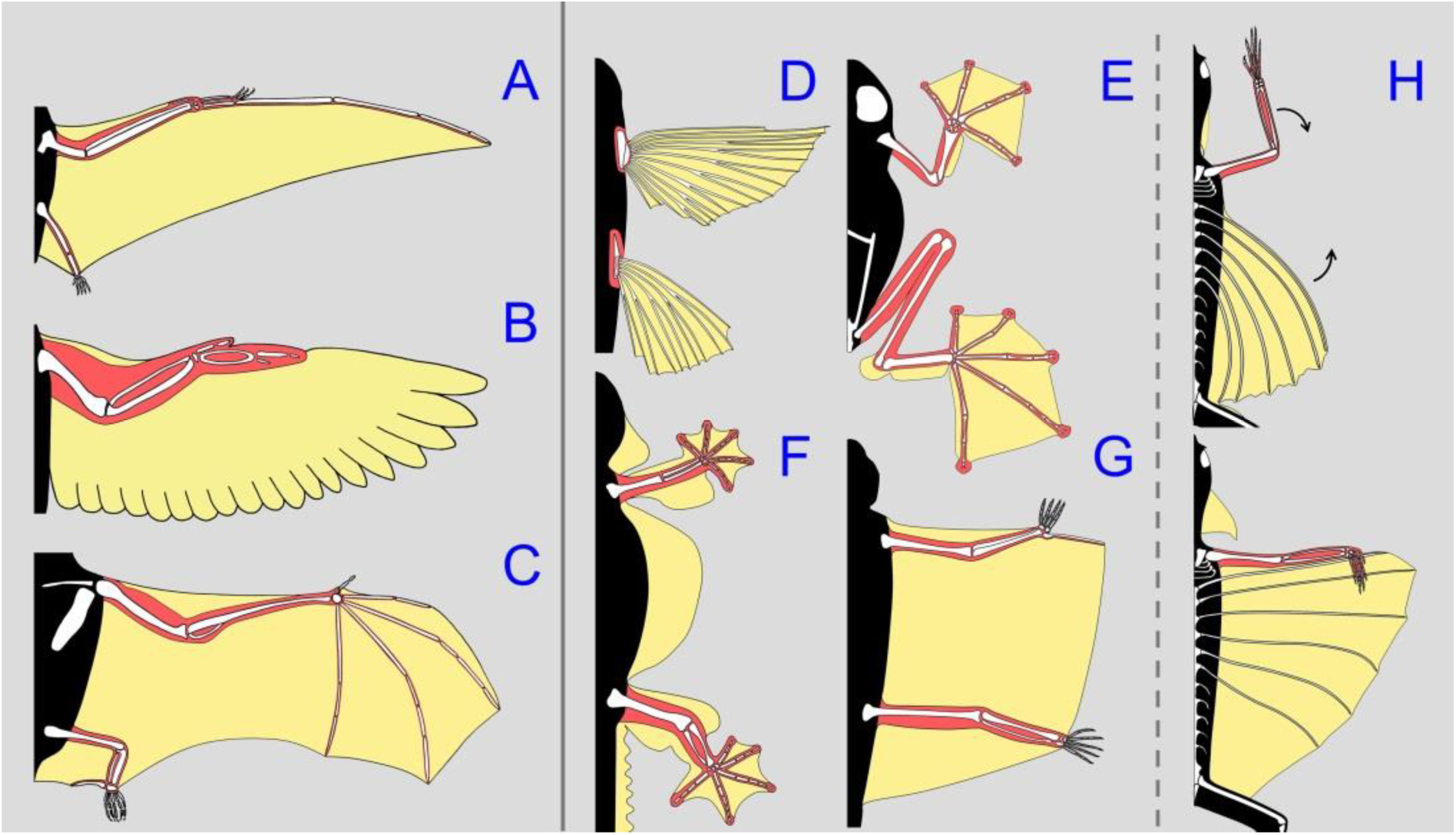
Wings and patagia of vertebrate groups with flapping (A–C) and gliding flight (D–H). Colours mark the major aerodynamic surfaces (yellow) and the skeletal and muscular structures that control them (red). A, pterosaur (*Rhamphorhynchus*, extinct); B, bird (*Columba*); C, bat (*Phyllostomus*); D, flying fish (*Hirundichthys*); E, flying frog (*Rhacophorus*); F, parachuting gecko (*Ptychozoon*); G, flying squirrel (*Petaurista*); H, flying lizard (*Draco*). In *Draco* lizards, the controlling unit is connected to the lift-generating surfaces only for the duration of the flight.

## DISCUSSION

My findings demonstrate that, contrary to previous assumptions, the forelimbs of *Draco* lizards are not held free next to the body during flight but constitute an essential part of the wing. This wing is unparalleled in the animal kingdom, as it represents the only case in which the lift-generating and the controlling units are formed independently by different parts of the body and must be connected to each other at the beginning of the gliding flight (Figs. 1–3). Apart from few groups of gliding or parachuting animals that use only their flattened bodies and unmodified, outstretched limbs to generate lift and drag forces (Yanoviak, Dudley & Kaspari, 2005; Vanhooydonck *et al.*, 2009; Socha *et al.*, 2015), all other groups of flying and gliding animals have developed enlarged aerodynamic surfaces, i.e. wings and patagia, that are permanently attached to the skeletal and muscular elements that control them (Norberg, 1990; Lingham-Soliar, 2015; Socha *et al.*, 2015). The enlarged aerodynamic surfaces of vertebrates are usually attached to modified limbs or fins (Fig. 3). In contrast to the hindlimbs, which possess moderate modifications, such as a lateral compression and a row of enlarged scales at the trailing edge of the thigh (Russell & Dijkstra, 2001; McGuire & Dudley, 2011), the forelimbs of *Draco* lizards lack modifications that would increase the surface. Such modifications could be expected if the forelimbs were held free next to the body and used to generate greater drag and lift forces, like in parachuting geckos and frogs (Emerson & Koehl, 1990; Young, Lee & Daley, 2002; Fig. 3). The major advantage of the composite wing of *Draco* is that the forelimbs keep their entire movement range and full functionality for agile climbing and running when they are not used as the controlling unit of the wing. Although this study reports the control of the patagium through the forelimbs only in *D. dussumieri*, the behaviour can be recognised in previously published photographs of gliding specimens of other *Draco* species (McGuire & Dudley, 2005; Socha *et al.*, 2015; Lee, 2015). Given the conserved patagial and forelimb morphology across all species of *Draco* (McGuire & Dudley, 2011; online supporting information, Table S1), the patagium is very likely controlled in the same way by all species of the genus.

The patagium of *Draco* differs functionally from the patagia of the parachuting geckos *Ptychozoon* and *Hemidactylus* (Fig. 3F), as the latter are unsupported by ribs, not controlled by muscles, and unfold passively as they catch air during descent (Russell & Dijkstra, 2001). Functionally, the patagium of *Draco* closely resembles the plagiopatagia of gliding squirrels and colugos, which extend between the arms and legs and are controlled by limb movements (Socha *et al.*, 2015; Fig. 3G). The patagium of *Draco* lizards, however, is not spread as a result of the limbs assuming a posture while airborne, but has to be deliberately grasped and extended.

The wing of *Draco* is characterized by distinct adaptive morphological features. According to aerodynamic theory, the camber of the aerofoil and the presence of a thick leading edge compared to the thin trailing edge create greater lift forces than flat wings could achieve (Norberg, 1990; Lingham-Soliar, 2015). The pronounced adduction of the wrist during the handling of the patagium enables the fingers to exert maximum forward traction on enlarged scale rows along the first two pairs of ribs on the dorsal surface of the patagium (Russell & Dijkstra, 2001). *Draco* species are able to deviate the wrist ulnarly, but not markedly radially, whereas other arboreal agamid lizards can neither adduct nor abduct their wrists (online supporting information, Table S1). Therefore, the adduction ability is obviously not related to climbing activities but appears to be a specific adaptation to grasp the patagium. As *Draco* lizards show a conserved morphology of the patagium in all species (McGuire & Dudley 2011) and the extended forelimb constantly reaches close to the lateral margin of the leading edge (online supporting information, Table S1), the need for control of the patagium through the forelimbs is probably an important constraint that prevents further rib elongation and increase in wing area.

The fact that a patagium can be controlled by largely unmodified limbs needs to be taken into consideration when interpreting finds of possible fossil gliders. A number of fossil lineages, including the Late Permian *Coelurosauravus*, the Late Triassic *Kuehneosaurus*, *Kuehneosuchus* and *Icarosaurus*, the Late Triassic *Mecistotrachelos*, and the Early Cretaceous *Xianglong*, possess elongated ribs or bony rib-like structures that are hypothesized to have supported a patagial membrane and thus resemble the glide-associated morphological modifications of the modern *Draco* lizards (Robinson, 1962; Colbert, 1970; Frey, Sues & Munk, 1997; Fraser *et al.*, 2007; Li *et al.*, 2007; Stein *et al.*, 2008). These fossil taxa are assumed to have glided through the air with the hands held free next to the body and to have changed direction by unilateral adjustments of the aerofoil through contractions of the trunk musculature (Colbert, 1970; Frey *et al.*, 1997; Stein *et al.*, 2008; McGuire & Dudley, 2011). Since *Draco* lizards use the forelimb to control the patagium, it is reasonable to presume that the fossil gliders regulated their glide path in a similar way. Skeletal properties of the fossil gliders allow this interpretation. The forelimb is shorter than the first elongated rib in these species and would have constituted a straight, thickened leading edge when extended and attached to the patagium. To hold on to the dorsal surface of the patagium, adduction of the wrist is advantageous, a condition which is apparent in the holotypes of *Icarosaurus siefkeri* and *Mecistotrachelos apeoros* and in a well-preserved specimen of *Coelurosauravus jaekeli* (Colbert, 1966; Frey *et al.*, 1997; Fraser *et al.*, 2007). Hence, it seems plausible that these early reptile gliders likewise controlled the patagium with their forelimbs. This would imply that the manner how the modern *Draco* lizards form and control an aerofoil while simultaneously retaining full movability of the forelimb was developed convergently in the past by several non-related reptile lineages.

## ACKNOWLEDGEMENTS

I am grateful to R. Rao, D. Bhaisare, A. Giri and colleagues at the Agumbe Rainforest Research Station for their help during my stay. W. Böhme (ZFMK) kindly provided access to material under his care.

## SUPPORTING INFORMATION

Table S1. Results of the morphological examination of voucher specimens of arboreal agamid lizards. For details and abbreviations see Materials and Methods. Symbols indicate the ability to deviate the wrist more than 80° (*) or less than 20° (–).

## REFERENCES

Colbert E. H. (1966) A gliding reptile from the Triassic of New Jersey. American Museum Novitates 2246: 1–23.

Colbert E. H. (1967) Adaptations for gliding in the lizard *Draco*. American Museum Novitates 2283: 1– 20.

Colbert E. H. (1970) The Triassic gliding reptile *Icarosaurus*. Bulletin of the American Museum of Natural History 143: 85–142.

Decramer M., Kelly S. & de Troyer A. (1986) Respiratory and postural changes in intercostal muscle length in supine dogs. Journal of Applied Physiology 60: 1686–1691.

Dudley R., Byrnes G., Yanoviak S. P., Borrell B., Brown R. M. & McGuire J.A. (2007) Gliding and the functional origins of flight: biomechanical novelty or necessity? Annual Review of Ecology, Evolution, and Systematics 38: 179–201.

Emerson S. B. & Koehl M. A. R. (1990) The interaction of behavioral and morphological change in the evolution of a novel locomotor type: “Flying” frogs. Evolution 44: 1931–1946.

Fraser N. C., Olsen P. E., Dooley A. C. & Ryan T. R. (2007) A new gliding tetrapod (Diapsida: ?Archosauromorpha) from the upper Triassic (Carnian) of Virginia. Journal of Vertebrate Paleontology 27: 261–265.

Frey E., Sues H. D. & Munk W. (1997) Gliding Mechanism in the Late Permian reptile *Coelurosauravus*. Science 275: 1450–1452.

Günther A. C. L. G. (1872) On the reptiles and amphibians of Borneo. Proceedings of the Zoological Society of London 1872: 586–600.

Hairstone N. G. (1957) Observations on the behavior of *Draco volans* in the Philippines. Copeia 1957: 262–265.

John K. O. (1967) Observations on the mating behaviour and copulation in *Draco dussumieri* Dum. & Bib. (Reptilia: Sauria). Journal of the Bombay Natural History Society 64: 112–115.

John K. O. (1970) On the ‘patagial musculature’ of the South Indian flying lizard *Draco dussumieri*, Dum & Bib. British Journal of Herpetology 4: 161–168.

Lee C. (2015) On the wings of dragons. Online article published 14 January 2015, available at http://www.wildborneo.com.my/blog/ (last accessed 8 November 2016).

Li P. P., Gao K. Q., Hou L. H. & Xu X. (2007) A gliding lizard from the early Cretaceous of China. Proceedings of the National Academy of Sciences U.S.A. 104: 5507–5509.

Lingham-Soliar T. (2015) The vertebrate integument, volume 2: Structure, design and function. Berlin & Heidelberg: Springer.

Maindron M. M. (1890) Dragons, fabled and real. Popular Science Monthly 36: 808–813.

Marsden W. (1811) The history of Sumatra, containing an account of the government, laws, customs and manners of the native inhabitants. Third edition. London: The author, J. M’Creery, Black-Court and Longman, Hurst, Rees, Orme, and Brown, Paternoster-Row.

McGuire J. A. (2003) Allometric prediction of locomotor performance: An example from Southeast Asian flying lizards. The American Naturalist 161: 337–349.

McGuire J. A. & Dudley R. (2005) The cost of living large: Comparative gliding performance in flying lizards (Agamidae: *Draco*). The American Naturalist 166: 93–106.

McGuire J. A. & Dudley R. (2011) The biology of gliding in flying lizards (genus *Draco*) and their fossil and extant analogs. Integrative and Comparative Biology 51: 983–990.

Mori A. & Hikida T. (1993) Natural history observations of the flying lizard, Draco volans sumatranus (Agamidae, Squamata) from Sarawak, Malaysia. Raffles Bulletin of Zoology 41: 83–94.

Norberg U. M. (1990) Vertebrate flight. Berlin: Springer.

Ratnovsky A., Elad D. & Halpern P. (2008) Mechanics of respiratory muscles. Respiratory Physiology & Neurobiology 163: 82–89.

Robinson P. L. (1962) Gliding lizards of the Upper Keuper of Great Britain. Proceedings of the Geological Society of London 1601: 137–146.

Russell A. P. & Dijkstra L. D. (2001) Patagial morphology of *Draco volans* (Reptilia: Agamidae) and the origin of glissant locomotion in flying dragons. Journal of Zoology 253: 457–471.

Seba A. (1734) Locupletissimi rerum naturalium thesauri accurata descriptio, et iconibus artificiosissimis expressio, per universam physices historiam. Cui, in hoc rerum genere, nullum par exstitit. Ex toto terrarum orbe collegit, digessit, descripsit, et depingendum curavit. Tomus I. Amstelaedami: J. Wetstenium, & Gul. Smith, & Janssonio-Waesbergios.

Socha J. J., Jafari F., Munk Y. & Byrnes G. (2015) How animals glide: from trajectory to morphology. Canadian Journal of Zoology 93: 901–924.

Stein K., Palmer C., Gill P. G. & Benton M. J. (2008) The aerodynamics of the British late Triassic Kuehneosauridae. Palaeontology 51: 967–981.

Vanhooydonck B., Meulepas G., Herrel A., Boistel R., Tafforeau P., Fernandez V. & Aerts P. (2009) Ecomorphological analysis of aerial performance in a non-specialized lacertid lizard, *Holaspis guentheri*. Journal of Experimental Biology 212: 2475–2482.

Whitelaw W. A., Ford G. T., Rimmer K. P. & de Troyer A. (1992) Intercostal muscles are used during rotation of the thorax in humans. Journal of Applied Physiology 72: 1940–1944.

Yanoviak S. P., Dudley R. & Kaspari M. (2005) Directed aerial descent in canopy ants. Nature 433:624– 626.

Young B. A., Lee C. E. & Daley K. M. (2002) On a flap and a foot: Aerial locomotion in the “Flying” Gecko, *Ptychozoon kuhli*. Journal of Herpetology 36: 412–418.

